# Mapping the Learning Curves of Deep Learning Networks

**DOI:** 10.1101/2024.07.01.601491

**Authors:** Yanru Jiang, Rick Dale

## Abstract

There is an important challenge in systematically interpreting the internal representations of deep neural networks. This study introduces a multi-dimensional quantification and visualization approach which can capture two temporal dimensions of a model learning experience: the “information processing trajectory” and the “developmental trajectory.” The former represents the influence of incoming signals on an agent’s decision-making, while the latter conceptualizes the gradual improvement in an agent’s performance throughout its lifespan. Tracking the learning curves of a DNN enables researchers to explicitly identify the model appropriateness of a given task, examine the properties of the underlying input signals, and assess the model’s alignment (or lack thereof) with human learning experiences. To illustrate the method, we conducted 750 runs of simulations on two temporal tasks: gesture detection and natural language processing (NLP) classification, showcasing its applicability across a spectrum of deep learning tasks. Based on the quantitative analysis of the learning curves across two distinct datasets, we have identified three insights gained from mapping these curves: *nonlinearity, pairwise comparisons*, and *domain distinctions*. We reflect on the theoretical implications of this method for cognitive processing, language models and multimodal representation.

**Author summary:** Deep learning networks, specifically recurrent neural networks (RNNs), are designed for processing incoming signals sequentially, making them intuitive computational systems for studying cognitive processing that involves dynamic contexts. There has been a tradition in the fields of machine learning and neuro-cognitive science to examine how a system (either humans or models) represents information through various computational and statistical techniques. Our study takes this one step further by devising a technique for examining the “learning curves” of deep learning networks utilizing the sequential representations as part of RNNs’ architectures. Just as humans develop learning curves when solving problems, the introduced method captures both how incoming signals help improve decision-making and how a system’s problem-solving abilities enhance when encountering the same situation multiple times throughout its lifespan. Our study selected two distinct tasks: gesture detection and emotion tweet classification, to illustrate the insights researchers can draw from mapping models’ learning curves. The proposed method hinted that gesture learning experiences are smoother, while language learning relies on sudden knowledge gains during processing, corroborating the findings from previous literature.

## Introduction

Over the past decade, deep learning and neural networks have achieved remarkable performance in prediction and classification tasks in various domains, from machine translation and object recognition, to autonomous driving and reinforcement learning [1, 2]. However, the increasing complexity of deep neural networks (DNNs) creates a challenge for interpreting these models. Researchers have expressed concerns that deep learning operates like a black box due to its impressive performance but poor explainability [3–6]. This lack of transparency can be problematic in both research and applied settings, especially if there is an over-reliance on such models for decision-making. Models can be biased in various ways, such as using stereotypes, and without understanding their underlying properties, they may fail to align with the goals of human designers [7, 8].

There is a long tradition in cognitive science and computational neuroscience of examining the internal representations of models [9–12]. One reason for this is to determine if a model’s features or processes reflect processes of the human mind. Such models can be informative for inferring properties of human mental processing, and so have direct theoretical implications. For example, Elman [13] and others [14] showed that recurrent neural networks (RNNs) can learn patterns sufficiently complex to resemble human grammar. By examining the internal activations of these recurrent networks, they showed that these systems are driven by graded, statistical features.

Words are not discrete “symbols” but scalar vectors conditioned by linguistic context in time [15]. This was taken to challenge theories that see language as a purely abstract and symbolic recursive process [16].

More recently, these questions have become prominent because of the success of large-scale models, especially DNNs [17] and large-language models [18]. For example, in the case of language models, examining the internal processes of the BERT Transformer-based architecture has shown that it may recapitulate common natural language processing (NLP) pipelines [19–21]. Recently, Chang and Bergen [22] found that the frequency and n-gram structure of word tokens significantly alters the training for language Transformer models learning these words.

Inspired by this prior work, in this paper, we devise a technique for examining the learning trajectories of deep learning models, in particular DNNs. There is historical precedent for our approach, too. McClelland, Rogers and others have studied the underlying knowledge of neural networks by tracking them as they learn [10, 23]. Despite this classic work in cognitive science, it is uncommon to see deep learning models that track progress as a way to unpack what is learned (e.g., going beyond simple RMSE curves; but see also [18], for a counterexample). We term this tracking a “learning curve,” as it resembles research on how human learners process incoming information and improve decision-making through iterations under different situations. A unique benefit with simulation is the possibility to examine many dimensions and measurements of the neural network over time. In the next section, we review recent work on interpreting DNNs and related models. We then introduce our approach based on learning curves.

## Background

To date, various techniques have been proposed to interpret DNNs. For example, many model interpretability techniques provide a task-specific and local explanation, such as saliency or attention maps that visualize the localized influence of a region on the output. These ad-hoc approaches are sometimes unstable as even changing a single pixel could substantially affect the local relationships between input signals and output data [24]. While most techniques emphasize the qualitative interpretation of a model, such as what has been learned or captured by the DNN, the lack of quantifiable measurements makes the cross-model comparison even more challenging [4, 5, 25].

In a foundational article on deep learning, LeCun and the colleagues [1] characterized DNNs as representation-learning methods utilizing multilayered large neural network-style models. Representations can be viewed as mental objects capturing semantic properties either observable or unobservable [26]. Different from traditional machine learning models, DNNs display remarkable flexibility and efficiency in encoding lower-level input signals, such as pixels, audio frequencies, or word tokens, into multidimensional vectors at a sophisticated level through multilayered nonlinear transformations [27]. These transformations generate multiple levels of representations that learn hierarchies of features at each layer [1]. The continuous numerical vectors (or hidden vectors) learned at each level are commonly referred to as “embeddings” and serve as dense representations of the original input data. Following this construction, numerous studies have demonstrated the correspondence between DNN-generated and real-world distributed representations among words and sentences [11, 28], speeches [29], images [9], objects [30] and scenes [31, 32]. Representational learning in DNNs offers fundamental contributions to cognitive science, as it can inform how cognitive systems process and organize knowledge, facilitating the comparison of learning processes between humans and machines [33–35].

Recently, several studies in cognitive science and neuroscience have highlighted the importance of integrative modeling between computation, human brains and behaviors [12]. Beyond comparisons of static end-point knowledge, scholars have begun exploring the potential correspondence in learning and information processing between DNNs and human cognitive systems, given that the current performance of DNNs can already approximate human performance across various domains [36–38] (see [18] for a review). For instance, the representations (i.e., embeddings) extracted from multilayered DNNs have shown significant accuracy in predicting neural and behavioral responses in humans throughout the hierarchy of learning and processing. This evidence spans multiple neural-behavioral measurements (e.g., fMRI, EEG, ECoG), modalities (e.g., visual, auditory, and language processing), and model architectures (ranging from simple embedding models like GloVe to more complex neural networks such as RNNs, convolutional neural networks, and transformer models (for further details, see [9, 11, 12, 23, 39, 40].

Given the extensive alignment observed between DNN embeddings and neural-behavioral activities, and their presumed meaningful representational mapping with human cognitive systems, an analysis of how they emerge in learning would seem important to understand these relationships. Goldstein and colleagues [11] identify the temporal correspondence between layer-by-layer embeddings in GPT-2 and evolving neural activities in language areas. However, this temporality is restricted to layerwise representations (from low-level to high-level representations) rather than how streams of signals have been received and processed by models or brains [38]. Our aim in this paper is to use temporal analysis in a systematic way by separating and tracking the time course of a network’s learning across classification tasks, thereby enhancing the understanding of the emergence of representations in DNNs.

### The current study

The goal of this study is to unpack the “learning curve” of DNNs through a sequence of hidden representations when the model encounters any temporal processing tasks across its training. To do so, we sample the network’s performance by using its embedding vectors to classify groups of items in its training input. This allows us to map out the progression of the network’s discriminations across these groups – how the network’s internal knowledge, in the form of embedding vectors, evolves during training.

The model architecture we focus on is RNNs due to their capacity to model sequential data and time-dependent tasks [41], such as text generation, speech recognition and stock market prediction. Although other deep learning architectures, such as convolutional neural networks and Transformers, also have the capacity to process time-series data, RNNs are explicitly designed for processing sequential data, as they effectively capture temporal dependencies through their recurrent connections. This mechanism makes RNNs an intuitive architecture for studying cognitive processing that involves dynamic, changing contexts [42].

In this study, we propose a generalizable interpretability approach that maps the global learning curves of DNNs based on changes in “predictability” or “information captured” between embeddings and output across time steps of the input data. To provide clarity, we use the term “learning curve” in this study to denote the holistic approach and intention behind mapping the underlying processing and developmental journey of DNNs. The method we propose separates two parts of the learning curve, one based on overall training, and another based on processing within input items during training. First, the “developmental trajectory” signifies the long-term learning process of DNNs, which is simulated by the increasing number of epochs (i.e., complete passes through an entire training dataset). Second, the “(information) processing trajectory” refers to the momentary accumulation of information across all timesteps (within an epoch), extractable from the performance of the RNN layer (see Fig 1 for a conceptual illustration). We detail each of these further below.

**Fig 1.**
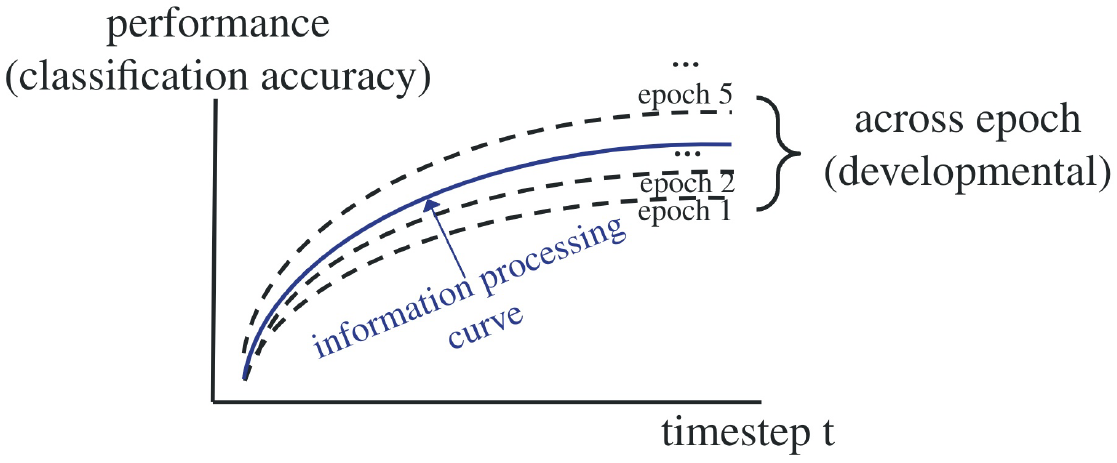
A conceptual illustration of learning curves in DNNs. The solid curve represents an individual information processing trajectory (across timesteps). For example, as an agent receives more signals over time in one session, its prediction of the gesture increases. The bundle of dotted lines represents a developmental trajectory (across epochs). For instance, this agent improves its ability to predict incoming gestures after repeatedly encountering similar patterns.

To illustrate the generalizability of our approach, we chose two distinct classification tasks: (i) sentence classification and (ii) gesture detection. These tasks differ widely in terms of their modalities. We predicted this would lead to variation in the underlying data generation processes and associated cognitive processing for each modality. In particular, gesture and body movements primarily result from the coordinated contraction and relaxation of muscles, with signals produced at later timesteps derived from the previous timesteps with relatively high autocorrelation [43]. On the other hand, verbal language, being a predominantly semantic modality, exhibits degrees of surprisal and arbitrariness that enhance the cognitive capacity of language processing [44–46]. Therefore, the expected developmental and information processing trajectories will likely exhibit distinguishable patterns across the two different tasks when the DNN system processes them respectively.

As we discuss in detail in later sections, this method has a few benefits. Through tracking the learning trajectory of a neural network, researchers can explicitly identify the appropriateness of a model for a given task as well as examine the properties of underlying input signals. This approach could also serve as a standalone visualization to map the accumulation of the underlying signals processed, which can facilitate research on multimodal neural networks and signal processing. Finally, mapping the learning curve of DNNs has the potential to assist future computational cognitive and neuroscience research and address whether the learning experiences of models also correspond to (or fail to correspond to) the temporal processing in human cognition in addition to the emphasis on static representations in the current literature.

In the following sections, we will provide a step-by-step method for visualizing the learning curves of neural networks, illustrate how to holistically interpret signal processing in them and quantitatively compare these curves across two different datasets.

## Methods

This research proposes a model-interpretability method that can extract the learning curve of sequence-based deep learning networks (e.g., RNNs). Inspired by cognitive science, the method measures the learning trajectory and underlying knowledge extracted by such networks. Tracking the learning curve of a neural network enables us to explicitly identify the model appropriateness of a given task, while also examining the properties of the underlying input signals. To illustrate the method, we use two temporal tasks: gesture detection and language processing as examples. This study therefore demonstrates that the method could inform a range of deep-learning tasks. In the next section, we introduce the model architecture of a general RNN-LSTM with fully connected layers, explain the nature of our datasets and the pipeline for using our approach in experimental simulations.

### Datasets

This study utilized two datasets: the Identity-free Video Dataset for Micro-Gesture Understanding and Emotion Analysis (iMiGUE) by [47] and the Emotion dataset from Hugging Face by [48] to examine the possibility of mapping learning curves for tasks involving temporality. The two temporal tasks (i.e., gesture detection and NLP) are distinct in terms of their modalities, lengths, and the steps required to extract and preprocess the features, thereby enhancing the diversity of data to illustrate the learning curve analysis. We detail the preparation of each dataset separately below.

### Emotion tweet classification

The Emotion dataset [48] consists of 20,000 English Twitter messages with six basic emotions (e.g., anger, fear, joy, love, sadness, and surprise) by adopting Plutchik’s [49] wheel of emotions, Ekman’s [50] six basic emotions, and hashtags in tweets. Tweets were annotated through noisy labels and distant supervision introduced by [51].

To prepare the emotion sentences for recognition by the RNN layer, we first applied the “basic English” tokenizer from torchtext to tokenize each tweet. Then, we used GloVe (Global Vectors for Word Representation), a word vectorization technique that does not rely on local word context statistics (local-context information), to vectorize each token in a sentence. GloVe was preferred over other token vectorization techniques like word2vec [52] due to its design to capture the universal meaning of each token/word, rather than the word’s meaning within a specific sentence or context. We opted for GloVe embeddings with 300-dimensional semantic features because it strikes a balance between capturing sufficient information and maintaining computational efficiency [35].

In theory, the input sequences of RNN are not required to have the same length. In practice, these sequences are padded or trimmed to the same length to optimize the computation in PyTorch. Accordingly, all tokenized tweets were padded to a consistent length of 66 tokens, which corresponds to the length of the longest tweet sample. The shape of each Emotion tweet follows a 300 dimensions × 66 timesteps (see Fig 2).

**Fig 2.**
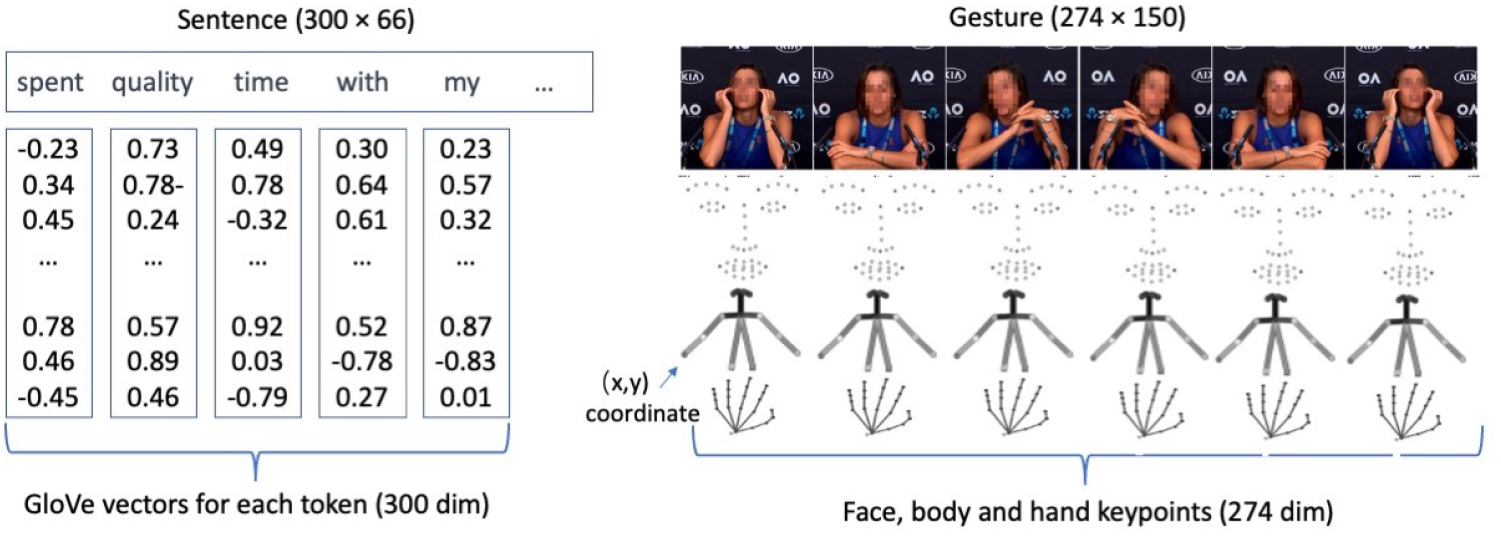
Sequential data for Emotion Tweet (left) and Gesture Classifications (right). NB: Gesture figures adapted from the iMiGUE dataset [47].

### Gesture classification

The iMiGUE is a high-quality dataset that contains 18,499 identity-free, ethnically-diverse, gender-balanced samples of 32 psychologically-meaningful micro-gestures (such as scratching an arm, adjusting the hair, or touching an ear [50]). These gestures were all curated from interview clips with athletes at post-match press conferences. Unlike other gesture and emotion datasets, which are typically drawn from staged performances or movie clips, the iMiGUE provides samples of actual gestures from real-life situations. This poses a more realistic, though more challenging, recognition task for deep learning networks [53, 54].

Due to copyright restrictions, the dataset includes only skeleton key points (rather than original interview clips) extracted from OpenPose, a multi-person computer-vision system that can simultaneously extract keypoints of the body, hands, face, and feet [55]. The advantages of using key point data are twofold. First, keypoint data conserves significant computing power; since each second of image sequences (i.e., matrices of pixels) may contain as many as 30 or 40 frames, running deep learning models on image sequences can be prohibitively expensive. Second, it provides better interpretability and understandability. Instead of being possible only through abstract information at the image-frame level, gesture detection can be operationalized as a sequential movement in keypoints across frames [56].

In total, 25 body, 70 facial, and 21 left hand and 21 right hand keypoints were extracted for each frame using OpenPose. Keypoint data were stored in the following format: [x0,y0,c0,x1,y1,c1…], in which (x, y) represents the coordinate of each keypoint and c indicates the confidence score of each keypoint-coordinate prediction. The confidence score was excluded in the following processing steps.

Although the gesture clips have just 39 frames on average, they have a higher standard deviation at 84 frames with a 75th percentile of 72 frames. We therefore set the padded length of the input sequence to 150 frames to ensure that our input data contained sufficient information for training a classifier. Thus, the input size for each gesture clip was 274 units (137 keypoints × 2) × 150 timesteps (see Fig 2).

It is important to note that our aim here is not to approach benchmark performance, but rather to examine successful learning. The learning curve analysis will show how that successful learning emerges, and which stimulus discriminations seem to underlie that emergence. Since even the state-of-the-art neural network can achieve only 55% accuracy on this multi-classification task [47], we further grouped the 32 micro-gestures into six categories (body, head, hand, body-head, head-hand movements and an absence of gestures) to ensure that our RNN-LSTM was indeed “learning” when we attempted to map its learning curve.

### Model architecture

We selected a model architecture that is standardized for sequential stimulus processing in the following way. First, we had a batch normalization layer, a primary RNN-LSTM layer, and then a fully connected layer to connect the final dense embeddings with the output classes (for classification). We offer details below.

### RNN-LSTM

RNN is capable of modeling sequential data and time-dependent tasks [41]. Its architecture represents an iterative function that takes an input sequence (*x*) and an internal state (*h*) from the previous timestep (*t - 1*) to predict the current timestep (*t*), Bthen updates the state as follows:

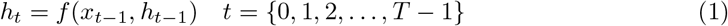

As the formula illustrates, each timestep *t* should theoretically reflect the information from 0 to *t* – *1*. We selected the RNN model for gesture detection because it can process temporal information under the assumption that the body movement in each timestep depends on signals in the previous timesteps.

While an RNN could leverage the context between elements by maintaining its internal state while processing the entire sequence, the “Vanilla RNN” layer experienced the vanishing-gradient problem during model training [57]. Therefore, long short-term memory (LSTM), which is represented by the function f in Equation 1, was introduced here as an additional state variable, called the cell state, for controlling specific information that needed to be kept or updated while processing the entire sequence [56]. Consequently, LSTM effectively reduced the vanishing-gradient problem encountered by RNN [57]. In general, LSTM is especially useful when the input data have a long temporal dependency, as was true of the gesture-recognition and sentence input data considered in this study.

### Feedforward network

The construction of our RNN-LSTM neural network followed the common practice in deep learning. In this network, a batch normalization was first applied to the data input, with the size batch size × timestep × input dimension (1 × 150 × 274 for gesture and 1 × 66 × 300 for NLP). The normalization was conducted across all timesteps of each data point to reduce variance within a data unit. Then, an RNN-LSTM was applied to the normalized input to convert temporal information to a dense embedding at each timestep. An RNN-LSTM with an equal hidden dimension was selected to simplify the tracking of embeddings in the later stage. Finally, a fully connected layer, without any activation, was applied to the embeddings in the last timestep t to predict output labels.

### Deep learning simulation

To ensure the generalizability of our learning curve mapping approach, we performed 15 rounds of simulation for both gesture and emotion detection tasks, each of which included 15 (6C2) pairwise binary classifications. We conducted 25 repetitions (reps) for each set of simulations (sims) to mitigate the idiosyncrasies of specific processing and developmental patterns we are extracting under each pairwise condition. In total, we collected 25 × 15 × 2 (reps × sims × tasks) = 750 runs of simulation data. Having an adequate number of simulation runs also enables us to observe clustering tendencies in the convergence and divergence of trajectory patterns across various tasks and different classes.

In each simulation, 20% of the shuffled samples were used as the test data, while the remaining data were further split into 80% training data and 20% validation data. While validation data were used for reporting each epoch’s model performance, test data were used for reporting the final model performance on unseen data.

The number of epochs indicates the number of times an entire dataset has been passed forward and backward through the neural network. As the number of epochs increased, the training performance shifted from underfitting to optimum to overfitting. The default number of epochs was specified as 20 to ensure that the models were learning from the data adequately.

Because deep learning is very sensitive to the unbalanced dataset, we applied a data augmentation to the training and validation dataset. Specifically, the minority class was up-sampled to match the number of data points in the majority class, which ensured a balanced dataset [58].

Default initiation from PyTorch was used to standardize the model specification across simulations. All simulation runs were trained using the Adam optimizer with a learning rate of 0.001, and the loss function used was the cross-entropy loss for all pairwise classifications.

Since deep learning is, computationally speaking, very expensive to train, the model was run on Nvidia RTX 3090 to expedite the processing. To maximize our training capacity, we used the mini batch, setting the batch size equal to 1, which increased the total amount of input data we could use for training.

To map a learning curve for each simulation, we first extracted embeddings of all LSTM timesteps from the hidden layer. We then applied multiple interpretable machine-learning models between these embeddings and the corresponding output labels to understand how the model’s confidence is updated during the LSTM timesteps.

### Embedding extraction

We extracted the LSTM array of all timesteps for all batches across all epochs on the test data to ensure that we were extracting embeddings from all developmental stages and the final model was adequately trained on the targeted task. Since we specified the same dimension of the input data for the LSTM hidden layer, for the gesture detection task, we obtained 150 LSTM arrays from 150 timesteps, each array having the size of 1 × hidden dimension (1 × 274). Similarly, we obtained 66 LSTM arrays in emotion tweet detection, each with a size of 1 × 300. We then stored the corresponding output labels of those arrays as an output array. Those LSTM arrays share one output array since they are embeddings at different timesteps of the same data points.

For binary classification, the labels were encoded as 0 and 1. Once the model had been trained and the embeddings extracted, we stored and processed those embeddings in CPU, which has a greater storing capacity than GPU.

### Mapping the learning curves

To approximate the processing curve of a model’s confidence across the LSTM timesteps, this study used a popular machine learning algorithm, k-nearest neighbors (KNN) for identifying embedding “separatables.” This algorithm classifies an object by using a majority vote of its neighbor data points [59, 60]. Because embeddings are seen as a high-dimensional physical (i.e., location-wise) projection of input data (as opposed to a multivariate representation [1]), distance-based models, such as KNN and support vector machine (SVM), are popular choices for examining DNN embeddings in previous studies. Specifically, we applied KNN to the data point (*x*_*t*_, *y*), where *x*_*t*_ is the LSTM embedding at *t* timestep and *y* is the corresponding output label and calculated the KNN accuracy across all timesteps (information processing trajectory) and all epochs (developmental trajectory), and thus captured the learning curve of the model.

### Quantifying performance across epochs

The foregoing sections describe how we setup and obtain binary classification performance of the RNN-LSTM across its training on two datasets. The main part of our proposed analysis here is to reconstruct and analyze these learning dynamics as they unfolded.

The constructed learning curves can be analyzed in many ways. In initial exploration, we tested various regression-based techniques. For example, we explored how the coefficients on a linear model that includes classification and epoch (acorss time) may be plotted and qualitatively assessed across data types. While these can partition and attribute the variance of these learning curves, regression coefficients across classifications are not easy to summarize. We instead chose simple summary statistics for network performance *across each trial*. These summary statistics can be tracked across epochs, and thus describe how the discriminability of the LSTM is evolving. We illustrate these summary statistics of an example trajectory below from a single run of a classification (Fig 3).

**Fig 3.**
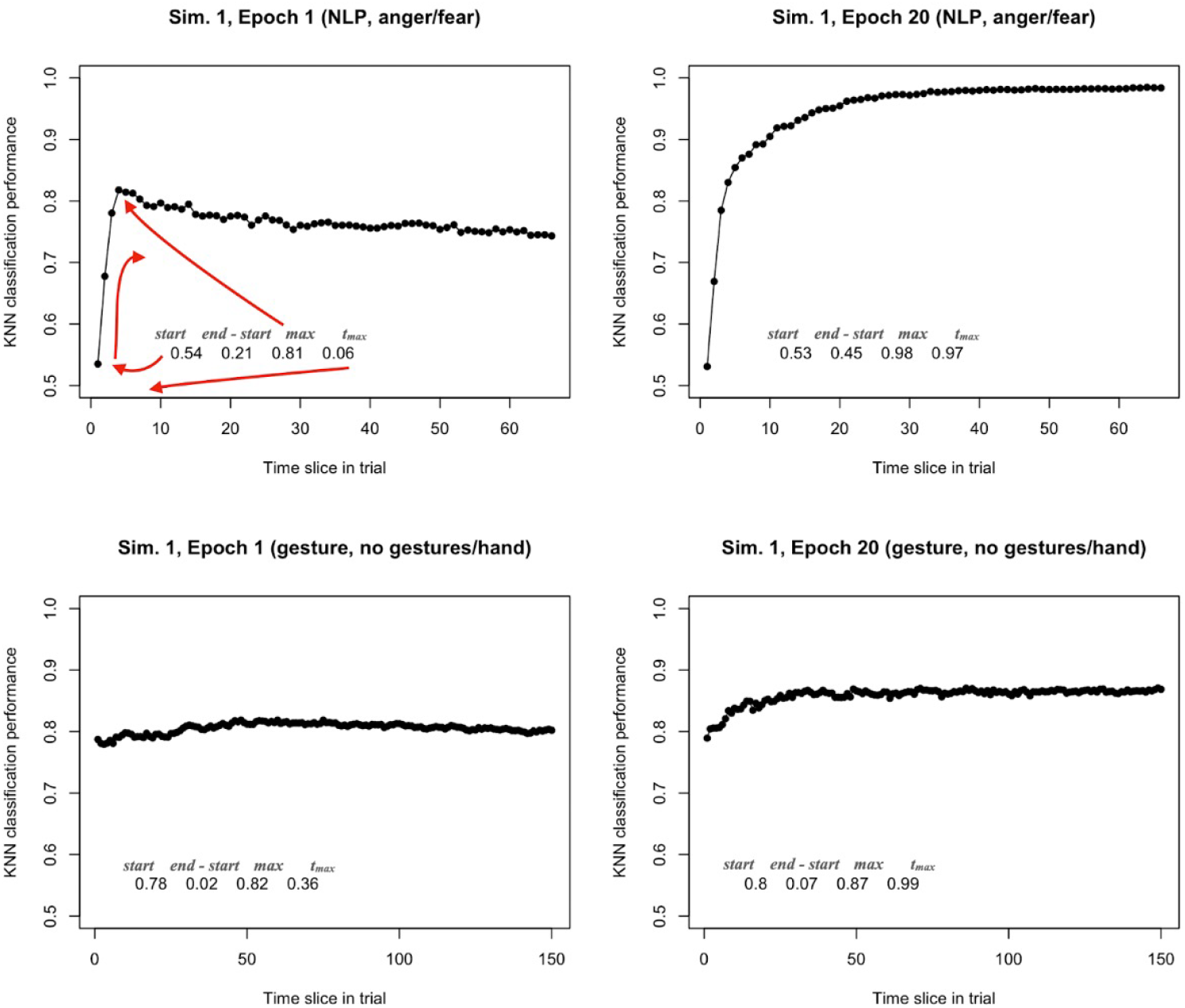
Examples at epoch 1 and 20 of binary classification across a test input for one run of the network. From epoch 1 to epoch 20, the LSTM’s embeddings are able to classify successfully over the test item, and we can characterize this success as a change to its performance using four metrics described in the main body of the text (*start, end - start, max, t*_*max*_). Top: Example item from the NLP task. Bottom: Example item from the gesture task.

As noted above, these learning curves show performance *within* an item presented to the network. We divide the text or gesture up into bins, and examine how LSTM’s embeddings can classify as the stimulus is incrementally presented to the network. The learning curve is plotted as proportion correct classification across the item. For each such curve, we extracted simple descriptives. First, we extract the maximum performance (*max*) and time bin at which that maximum was achieved in the presented item (*t*_*max*_). We find that *max* by itself is not sufficient to determine performance, especially earlier in training. This is because the network may exhibit unstable performance, dropping across subsequent time slices. This may be indicative of nonlinear learning shown in related domains [61]. This also suggests the network has learned something about the *initial* segments of an item, but the later segments wash out its performance as it has not yet encoded these later features. To capture this trend, we use *t*_*max*_ to assess when that maximum was achieved inside the item – the time slice of the observed maximum performance. In Fig 3 above, we show an illustration of this in the top left. The network achieved a performance of 0.81 on this particular item, but failed to sustain this performance as it dropped to near 0.75 as the sequence unfolds (at epoch 1). In the top right, the performance improves approximately monotonically across time slices (by epoch 20), and *t*_*max*_ is achieved near the final bin of the training item.

We also assess the performance at the start and end of the presented stimulus (*start, end*). These have distinct interpretations. Performance at the start may indicate the relative gains that can be expected from a stimulus item. As shown in the bottom panels of Fig 3, the gesture input already has performance above chance (∼ 0.80) after the very first segment of the stimulus item. This suggests the model rapidly exploits spatial information in gesture. A model that achieves near-perfect performance at start and sustains it does not need to be exposed to the subsequent stimulus. Indeed, a fourth and final measure we use is the subtraction of *end - start* performance. A high value on this measure suggests the network gets substantial information gains across a test item. For example, even at epoch 20 in the bottom right, the *end - start* of the gesture item is substantially lower than the simulation trained on the NLP task.

These measures can be plotted across epochs. Each learning curve now indicates how an item is being processed across an LSTM’s overall training. The measures are relevant to two timescales in the network’s behavior: developmental (or learning) and information-processing timescales. For example, across epochs, movement along the start measure represents how the network performs when it is presented with the item’s first time slice. Across epochs, *end - start* can describe the relative information gains from the item’s total presentation. If *t*_*max*_ is low, it indicates that the network’s performance may be deterred by subsequent time steps, suggesting more training is needed. Finally, *max* is simply the overall performance, related to the commonly used measures of mean error or performance.

We took the output from the KNN classification in Python, described above, and designed a sequence of R scripts to measure, visualize and quantify the trends in these four measures. R’s suite of visualization tools provided a convenient arena within which to view trends across epochs, and in these analysis scripts we also built linear models to statistically test the significance of these trends. All of the scripts in our methods are available at GitHub here: https://github.com/JoyceJiang73/Learning-Curves/. The R scripts only require input of simulation CSV data that contain as fields: binary classification labels, time slice, epoch, and performance measure (e.g., KNN classification performance).

## Results

### Start performance

In Fig 4, we show the LSTM performance at the *start* of an item across epochs of training. In general, the NLP dataset shows low, near-chance performance, while gesture classifications are already well above chance performance. This chance-level performance for NLP items stays consistent across the entire training period, though the gestural dataset shows some improvement. For the gesture dataset, this suggests that the network has some information about a classification before much of a training item is even shown to the network. It would indicate that gestural data has spatial information in the point coordinates of the body and is exploited by the network at the very first time bin. With language, it takes time for word embedding vectors to be integrated in the network.

**Fig 4.**
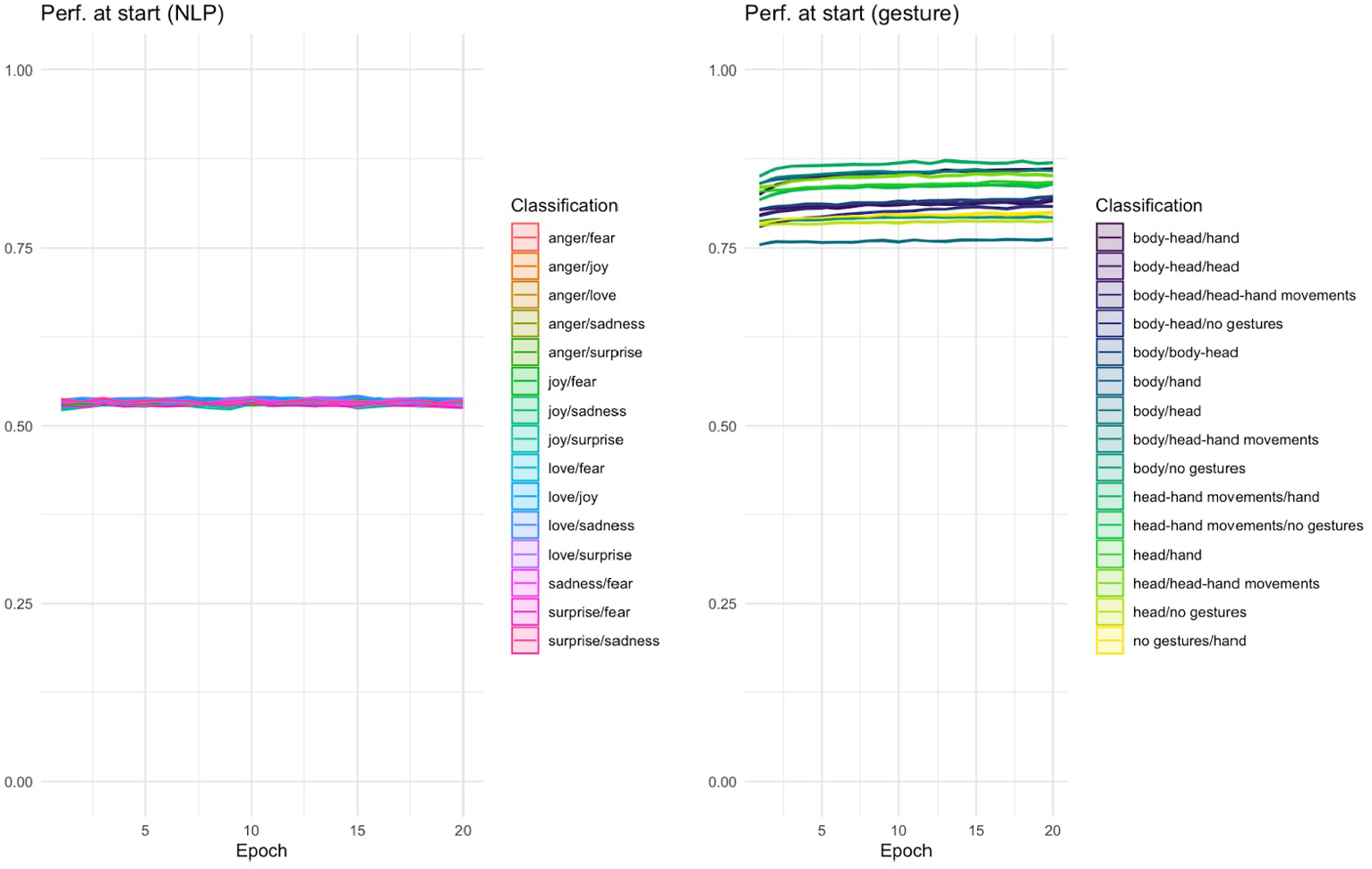
*Start* performance for pairwise classifications. *Start* metric over epochs, indicating that in the NLP task, the classification at the first time slice remains stable near change, whereas the LSTM’s embeddings for the gesture task are distinct across stimulus types, and also show slightly more improvement over training (while already being well above change relative to NLP).

To confirm these trends, we tested a linear model that predicted start performance by training data, showing that dataset accounted for about 98% of the variance seen in Fig 4 (*R*^2^ = 0.98, *p* < .00001). Moreover the tendency for the gestural training to show slight rise in learning whereas the NLP dataset is flat is expressed by a significant interaction when added to this model (*p* < .00001). The classification of the gesture data accounts for 95% of variance internally to that dataset (*p* < .00001), whereas for NLP classifications accounts only for 9.8% (lower but also significant, *p* < .00001). This would be predicted if the first word of an NLP task has low diagnostic accuracy for a classification, but the spatial variance over gestural classifications is much more informative.

### Max performance

Curiously, if one only investigated maximum performance across training, these classification tasks could be regarded as relatively similar in their behavior. As shown in Fig 5, both NLP and gesture datasets yield a classification performance that is high, between 0.75 and 1.0 depending on the classification. In the gesture dataset, there are more “difficult” classifications, shown by outliers in *max* performance across training. This can be helpful in diagnosing representational challenges in the network’s training, marking what pairs of training items may be more difficult to distinguish than others.

**Fig 5.**
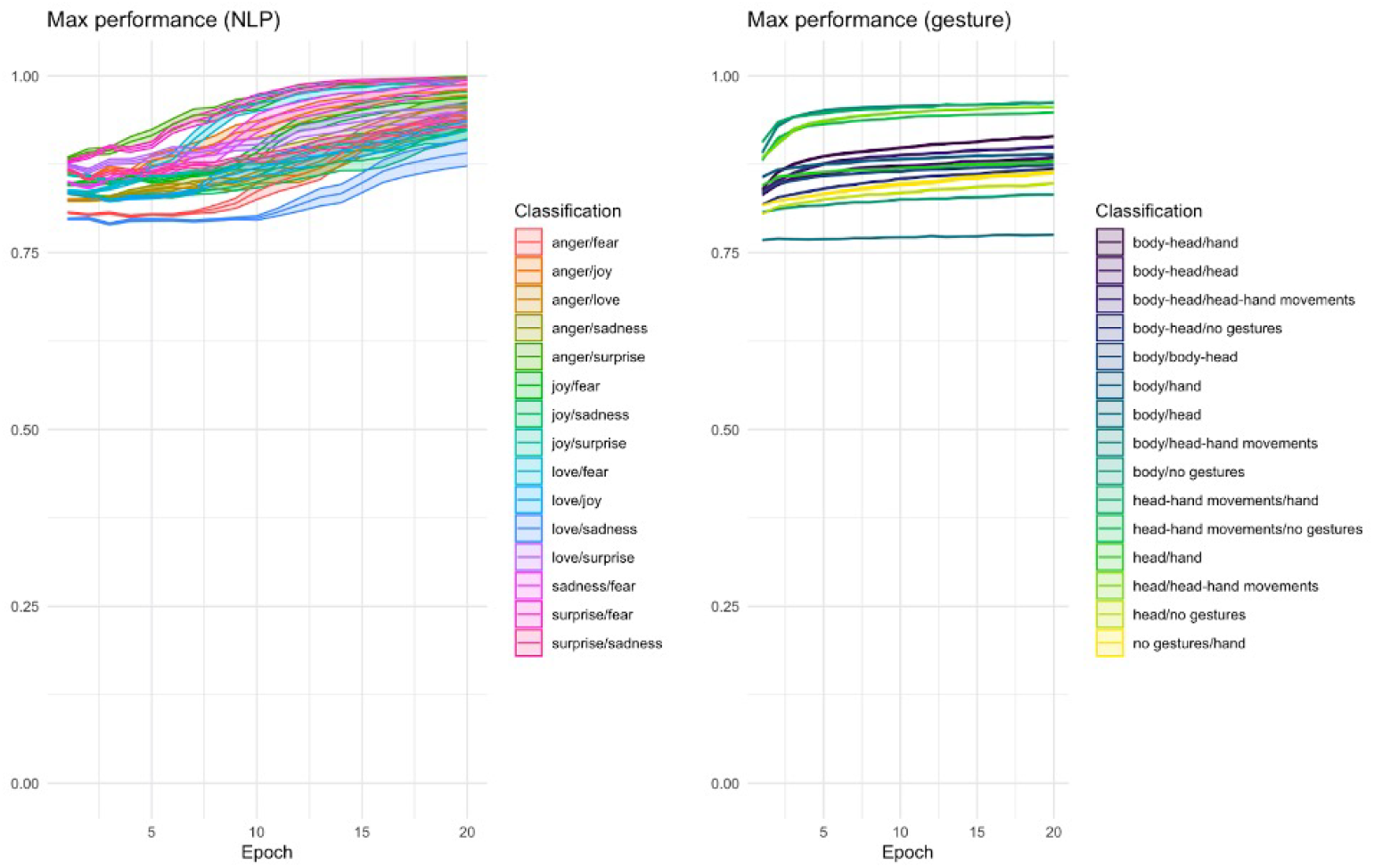
*Max* performance for pairwise classifications. Both NLP and gesture tasks show an increase in *max* performance over training epochs. On average this asymptotes near perfect performance but can vary depending on binary classification.

Again, as with the *start* measure, the *max* performance shows greater variance associated with classifications in the gesture case (92.6%, *p* < .00001) than the NLP case (26.7%, *p* < .00001). The difference between these two datasets is not as pronounced as in the *start* measure, as a linear model shows that only 2.4% of the variance is associated with the dataset in a linear model predicting max performance observed within a trial. Despite the small value, it is nevertheless significant and driven by the relatively higher performance in NLP (*p* < .00001).

### Time at max

We would expect that completed training in the LSTM should show that maximum performance should appear near the *end* of an item, as this would suggest that the network has extracted useful information in the whole presentation. Indeed, this is indicated by lower time-at-max values earlier in the training. During the first few epochs, the maximum value occurs proportionally earlier in a training item, similar to the example shown above in Fig 3. However in both NLP and gesture datasets, networks slowly extract features across the stimulus items for classification, as shown in Fig 6. The time gradually rises for all classifications, though it can also vary widely and tends to be more irregular in gesture.

**Fig 6.**
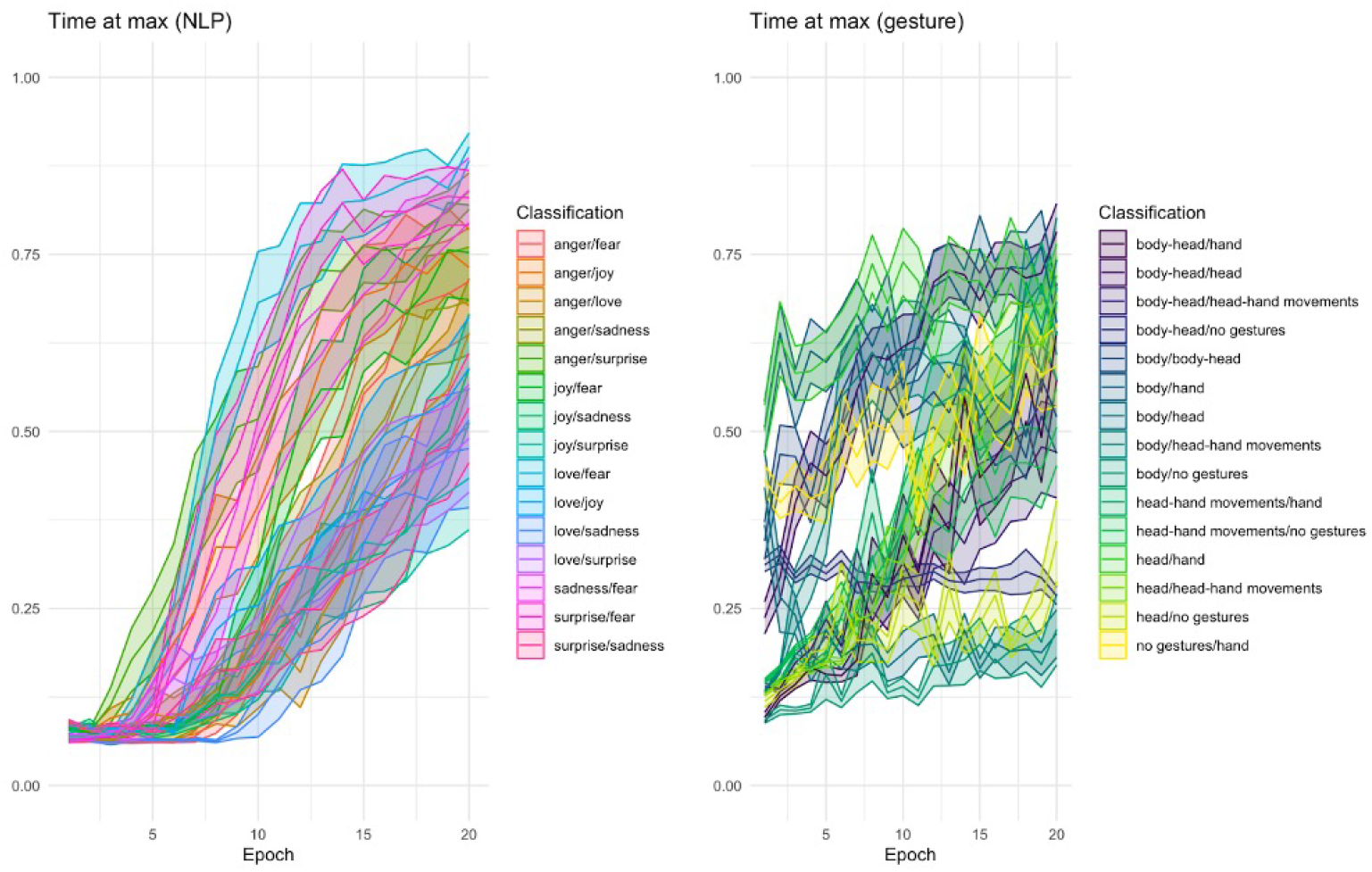
Time at max (*t*_*max*_) for pairwise classifications. Both NLP and gesture tasks show an increase in the proportion of time at which maximum performance is observed. Indicative of information gain, this rises in both – suggesting the LSTM comes to better integrate the full test items for its performance. However this is much more orderly in the NLP task than the gesture task.

In a linear model predicting *t*_*max*_ from data source, NLP and gesture are only slightly different, with 1.0% of the variance accounted for and NLP predicted to be slightly lower by about 0.06 proportionally (duration-scaled range 0 to 1). Despite this small effect, it is significant (*p* < .00001). Classifications in NLP and gesture both relate significantly in a linear model predicting time at max, with NLP classifications accounting for 10.0% of the variance and gesture classifications 30.3% (*p*’s < .00001).

### End - start performance

Finally, we wish to get a sense of how the LSTM models are extracting information across the stimulus item. One way to do this is to discern how much higher its performance is at the end of an item compared to its performance initially. This is where we see the greatest difference between the datasets. As shown as Fig 7, NLP shows substantial information gain across stimulus items. By the end of training, the start and end of trials involves considerable difference, and the positive value of this difference shows that the network improves its performance significantly across presentations. With gesture, the situation is quite distinct. In most classifications, there is indeed a rise. But because gesture is informative even at the *first* time bin of an item, there isn’t much information gain remaining. Indeed even the highest instances of this value hover near 0.10. So while performance on gesture is higher at first, the network may find the task more difficult in integrating that spatial information over time to improve performance. Indeed, one classification (body/head) shows information loss at subsequent time bins across simulations.

**Fig 7.**
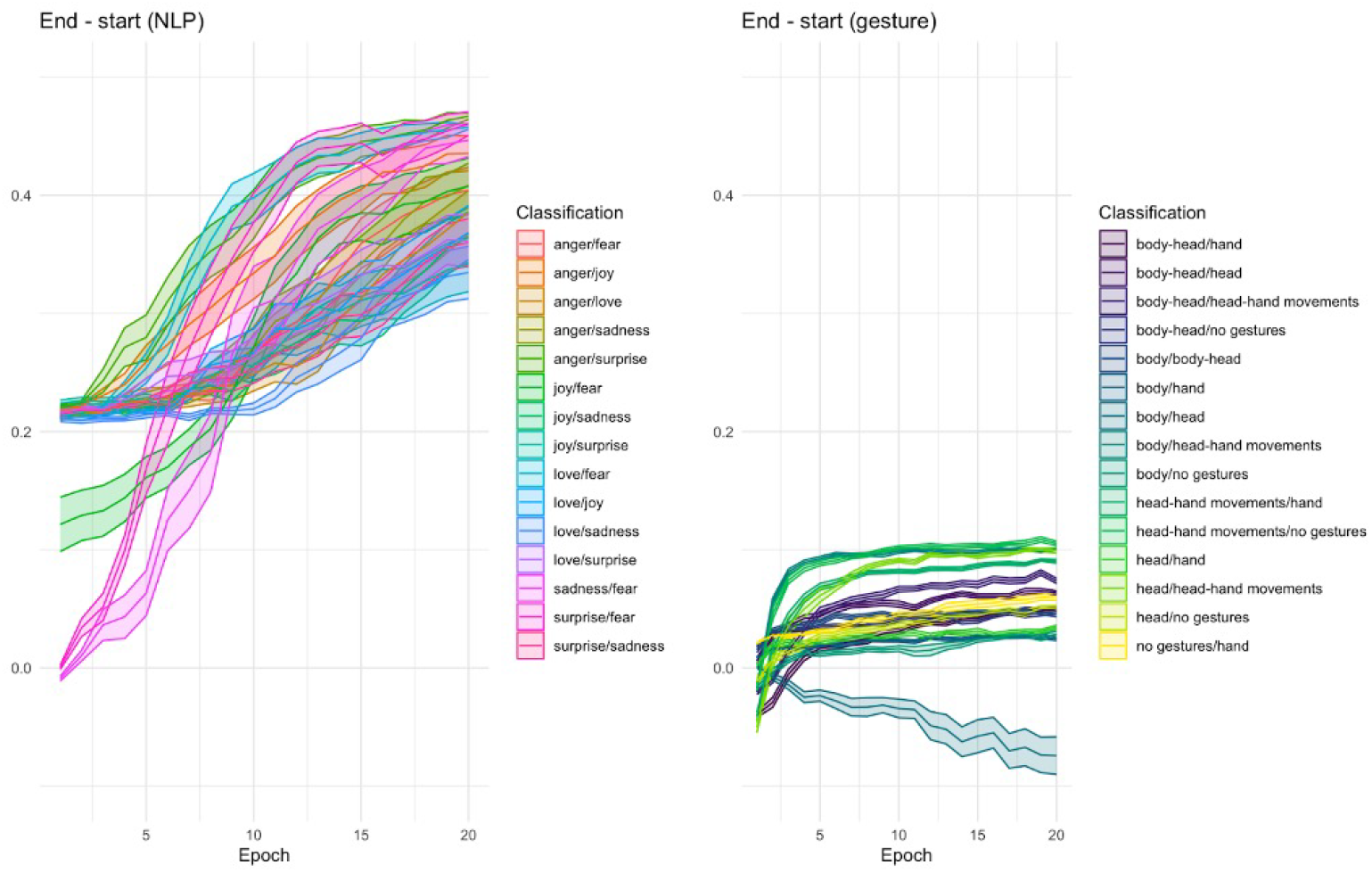
*End - start* performance for pairwise classifications. The information gain from start to end of stimulus item shows a stark difference between datasets. The NLP task shows that the LSTM’s performance rises as it integrates data from start to end. With gesture, the story is more complicated, showing less of a rise and in one case information loss in one classification.

As in the *start* measure, the difference between datasets is quite large. When data source is used to predict the *end - start* measure, 65% of the observed variance in this measure can be associated with data source (*p* < .00001), with NLP associated with much higher information gain than gesture results (*B* = 0.25). When looking at each data source separately, only 8.0% of the measure is associated with the classification pairs in NLP. In the gesture dataset, this association is 57.6%, showing much higher contribution of the class differences in gesture. Indeed, inspecting an interaction term between epoch and classification, as Fig 7 recommends, we find that a linear model with epoch and classification accounts for 74.4% in the case of gesture and only 51% in NLP (both *p*’s < .00001).

## Discussion

Recognizing the importance and challenges involved in systematically unpacking the internal representations of DNNs, this study introduced a multi-dimensional quantification and visualization approach, “learning curves,” which can capture two temporal dimensions of a model learning experience. First, it captures the “information processing trajectory,” how the network is doing as it processes test items. Second, it captures the “developmental trajectory,” describing how this processing is changing over training epochs. The former represents the influence of incoming signals on an agent’s decision-making, which is operationalized by the timestep within a single epoch of an RNN. The latter conceptualizes the gradual improvement in an agent’s decision-making abilities throughout its lifespan, operationalized by the iteration of epochs. The learning curve approach we illustrate in our two datasets shows that we can quantify and qualitatively investigate both of these dynamics within the same analysis.

Beyond the visual presentation and qualitative inspection of the multi-dimensional learning curves, this study further defines four measures: start performance (*start*, the initial capacity of each information processing), max performance (*max*, the maximum performance of each information processing), time at max (*t*_*max*_, when the current information processing reaches the maximum performance), and end – start performance (*end – start*, the overall performance gain in this information processing) – to facilitate quantitative comparisons across tasks and datasets. Based on the analysis of these four measures derived from the learning curves across two distinct datasets, we highlight three insights gained from mapping these curves: *nonlinearity, pairwise comparisons*, and *domain distinctions*.

### Nonlinearity

First, we observed nonlinearity in the learning experiences of DNNs, a characteristic recognized in previous literature as a key advantage for their advancement in addressing the most challenging tasks. Specifically, we found that RNNs exhibit different preferences for early versus late cues when addressing various sequential tasks. For instance, in the NLP task, we noted a more pronounced performance improvement occurring in the later stages of information processing (i.e., model development). In contrast, gesture learning tends to show quicker progress, with more variability across epochs and repetitions of simulations, suggesting that DNNs tend to rely on shortcuts, such as naive cues related to keypoint coordinates, for gesture classification, rather than focusing on high-level movement sequences. This shortcut-based “learning” also is evident in the higher initial performance (*start*) for gesture classification (*>* 0.75). On the other hand, NLP classification begins at around 0.50 (the at-chance probability) and exhibits greater performance gains in later epochs, indicating that the models classify based on high-level semantic sequences.

### Pairwise comparisons

Additionally, although multiclassification tends to exhibit collective model performance, our between-class pairwise comparisons reveal the presence of outliers within multiclassification. For example, “joy/fear”, “surprise/fear”, and “sadness/fear” demonstrate higher information gains across epochs compared to other emotion pairs, suggesting that these classes are further apart from each other. The “body/head” classification appears to experience learning challenges, possibly because these two movements have difficulty being completely separated, as the head’s movement in naturalistic data may inevitably coincide with that of the body due to joint coordination.

The combination of learning curve mapping and instance measures thus serves as an effective approach for “auditing” representations in multiclassification problems. The proposed pipeline improves model explainability beyond holistic evaluation of classification performance and ad-hoc attention visualization by unpacking pairwise class learning patterns to reveal any pairs that are unsuccessful in being discriminated or involve delayed knowledge gain. This granular examination allows modelers to better investigate a model’s appropriateness for the underlying task, as well as the properties of the processed input signals, reaffirming the value of our learning curve conceptualization.

### Domain distinctions

With these findings, it is tempting to infer that there are domain distinctions among different modality classification tasks. Gesture learning, for instance, may rely on smoother autocorrelated signals, potentially emphasizing spatial semantics as early cues, while language learning relies on higher degrees of surprisal, irregularities and arbitrariness, as suggested by previous literature (see [43–46]. These domain distinctions are valuable to cognitive science in understanding how humans process and distinguish between various modes of communication, shedding light on the neural mechanisms underlying the flexibility and adaptability of the human mind when processing different forms of information and communication modalities.

Though it is intuitive and tempting to explain these domain distinctions here, we cannot yet assert that these trends would hold for *all* NLP or gesture (keypoint) classification tasks, only the ones we investigate here. However the learning curve results would seem to align with an intuition of how linguistic symbols would be sequentially integrated into a neural network in contrast to the highly auto-correlated spatial information contained in gesture performance. Still, we cannot infer a broad generalization about “language vs. gesture” and only leave it as a potential path for future investigation. Indeed, this may be an additional benefit of a method like the one we present here. Learning curves could finely ascertain these domain distinctions, and expanding the set of data to test may permit generalization in future work. This may have theoretical implications itself. The learning curve analysis may provide information about the distinct sources of information from varied modalities. When neural models (or human brains, presumably) integrate distinctive sources of information, they may strengthen understanding of complex multimodal data by leveraging their unique information-processing and developmental benefits.

## Conclusion

Taken together, the temporal mapping (information processing and developmental trajectories) helps modelers more comprehensively understand the underlying learning and decision-making processes of a complex model architecture without delving into the intricacies of interpreting its internal representations directly [23, 38]. This kind of systematic and quantitative approach has recently gained popularity in both computational cognitive science and deep learning communities [5, 38, 62–64], as it facilitates multidimensional comparisons across models and modalities, which were previously seen as challenging for DNN-like models. The current study illustrates multiple techniques for analyzing model learning experiences and highlights three insights across different communication modalities based on these analyses. Future studies can utilize this learning curve mapping approach to enhance model interpretability studies by evaluating a model’s appropriateness for the task at hand, examining the properties of the underlying input signals, and assessing the model’s alignment (or lack thereof) with human learning experiences, which is also a critical consideration for computational cognitive science and neuroscience research.

## Acknowledgments

We thank the organizers, panelists, and audience for allowing us to present the initial version of our work at the 53rd Annual Meeting of the Society for Computation in Psychology (SCiP) and for sponsoring the registration fee. We are also grateful to the reviewers from the 45th Annual Meeting of the Cognitive Science Society for their comprehensive review and invaluable feedback. Additionally, we thank Hongjing Lu, Jungseock Joo, and Elisa Kreiss for their insightful feedback on the theoretical framework and methodologies.

## Author Contributions

**Conceptualization:** Yanru Jiang, Rick Dale.

**Data curation**: Yanru Jiang.

**Formal analysis**: Yanru Jiang, Rick Dale.

**Investigation**: Yanru Jiang, Rick Dale.

**Methodology**: Yanru Jiang, Rick Dale.

**Project administration**: Yanru Jiang, Rick Dale.

**Resources**: Yanru Jiang, Rick Dale.

**Supervision**: Rick Dale.

**Visualization**: Rick Dale, Yanru Jiang.

**Writing – original draft**: Yanru Jiang, Rick Dale.

**Writing – review & editing**: Yanru Jiang, Rick Dale.

